# *Escherichia coli* robustly expresses ATP synthase at growth rate maximizing concentrations

**DOI:** 10.1101/2021.02.17.431610

**Authors:** Iraes Rabbers, Frank J Bruggeman

## Abstract

Improved protein expression is an important evolutionary adaptation of bacteria. A key question is whether evolution has led to optimal protein expression that maximizes immediate growth rate (short-term fitness) across conditions. Alternatively, fitter genetic variants could display suboptimal short-term fitness, because they cannot do better or because they strive for long-term fitness maximization by, for instance, anticipating future conditions. To answer this question, we focus on the ATP-producing enzyme F_1_F_0_ H^+^-ATPase, which is an abundant enzyme and ubiquitously expressed across conditions. We tested the optimality of H^+^-ATPase expression in *Escherichia coli* across 27 different nutrient conditions. In all tested conditions, wild-type *E. coli* expresses its H^+^- ATPase remarkably close to optimal concentrations that maximize immediate growth rate. This work indicates that bacteria can achieve robust optimal protein expression for immediate growth-rate.

## Introduction

According to the accepted view on fitness, the fittest microorganisms produce most viable offspring over a given period of time and (dynamic) conditions.^1,2^ One long-term fitness strategy is to always aim for maximization of immediate growth rate (short-term fitness). Another strategy is to prepare for future conditions, leading to a suboptimal short-term fitness, but an optimal long-term fitness; for instance, by ensuring that adaptation times are generally short or by anticipatory expression of stress proteins.^3–6^ Which strategy is preferred depends on the evolutionary history of the organism^7,8^ and, likely, also the optimization capabilities of the regulatory circuitry of cells.^9^

Fitter microbes express proteins at better adapted concentrations or ones with better adapted (kinetic) properties. In principle, a cell can enhance its net biosynthetic rates, and hereby its immediate growth rate (short-term fitness), by increasing concentrations of needed metabolic enzymes – since the rate of an enzyme is generally proportional to its concentration.^10^ However, bounds exist on the total allowable protein concentrations in a cell.^11,12^ For instance, because proteins occupy space and finite transcriptional and translation resources limit biosynthetic processes.^11,13–15^ In addition, enzymes have per capita finite activities.^10^ Thus, the immediate growth rate of a microorganism has a maximum. Bacteria therefore need to solve a constrained “optimization” problem to optimally allocate their limited biosynthetic resources over all (growth-rate supporting) proteins in the cell.^9,11,13,16^ Examples of limited biosynthetic resources are, for instance, RNA polymerases, ribonucleic acids, ribosomes, amino acids, free-energy carriers, membrane space, and cytosolic space.^11,16^

That biosynthetic resources are limited in a bacterial cells is indicated by the fact that the expression of unneeded proteins reduces immediate growth rate, as their synthesis reduces concentrations of needed growth-supporting proteins.^12,16–18^ This was, for instance, illustrated by expressing beta-galactosidase during growth on a minimal medium supplemented with glucose rather than lactose.^12^

Tuning the concentrations of needed proteins (e.g. for growth or stress relief) to optimal levels for the current environment can maximize immediate growth rate, as then no biosynthetic resources are wasted.^19–22^ Alternatively, a needed protein can be over- or underexpressed, relative to its optimal level, because the organism cannot perform better or it is preparing itself for future conditions.^4,23^ In the latter cases, the expression level of the needed protein is suboptimal for immediate growth rate.

An optimal level appears to exists for any needed protein. Imagine that we slowly increase the concentration of such an enzyme. When its concentration is increased, starting from zero, the growth rate increases from zero too. The growth rate continues to rise as long as the enzyme is “underexpressed”.^19–22^ Eventually, the growth rate reduces when the enzyme is increased even further, i.e. when it is “overexpressed”.^12,19–22^ In between the underexpression and overexpression levels the growth rate reaches its maximum at the optimal expression level.^19–22^ In the overexpression regime, the growth rate reduces because too many limited biosynthetic resources are allocated to an enzyme that is no longer growth limiting.^12,17,18^ Prevention of protein overexpression is therefore a requirement for maximization of the immediate growth rate (short-term fitness). Since expression costs generally rise linearly with the expression level and the benefits maximize at a certain expression level (when other proteins become growth limiting), an optimum is found at intermediate expression levels where the difference between protein benefit and cost are maximal.^19,24^

Optimal expression levels have been found experimentally for a number of different proteins across microorganisms. For instance, for glycolytic proteins in *L. lactis*^25–27^; enzymes of the PTS system of *S. typhimurium*^28^; citrate synthase^29^, catabolic genes regulated by CRP^6^, H^+^-ATPase^30–32^, and the lac operon of *E. coli*^19^; and dozens of proteins of *S. cerevisiae*, involved in various processes.^20^ These studies provide strong evidence for protein expression optimization by microorganisms. Natural selection for growth rate at constant conditions, i.e. for immediate growth rate, should also in principle lead to evolution of protein expression towards its optimum.^16^ Also, this has been shown experimentally.^19,33^

Whether a microorganism expresses its proteins to an optimal level to maximize growth rate is nontrivial, especially if it is subjected to a dynamic environment. It could well be that an organism has adapted its expression level to circumstances it most often encounters (e.g. glucose as carbon source), but performs suboptimally in more unusual circumstances (e.g. mixed carbon/nitrogen sources). Moreover, the exact regulatory circuits remain unknown for many proteins and pathways, which hampers the prediction of the extent to which they are able to robustly tune expression levels throughout different environments. Experimental studies are therefore needed to assess optimality of protein expression. Studies so far have focused on optimal expression of proteins in only a few environments. Therefore, the extend to which *E. coli* is able to *robustly* optimize the expression level of a key protein through phenotypic adaptation (on short timescales, excluding adaptation through mutation and selection); thus, over a diverse set of different environments. This is the focus of this work.

## Results

### Optimal expression of F_1_F_0_ H^+^-ATPase across conditions

We chose to study the optimality of the expression of F_1_F_0_ H^+^-ATPase (ATPase). It catalyzes ATP synthesis, a vital cellular process. This membrane-embedded protein is composed of 8 subunits, each expressed from a single operon, i.e. *atpIBEFHAGDC*, with a single promoter.^34–36^ ATPase has a total mass of ∼545 kDa, making it 10 times as large as the average protein (54 kDa). Its size is 20% of the size of the largest protein, the ribosome (2700 kDa). ATPase generates vast amounts of ATP under aerobic conditions at the expense of the proton motive force that is maintained by an electron transport chain.^35^ While glycolysis alone generates ∼4 ATP molecules per glucose molecule, the activity of ATPase adds to that 28-38 ATP molecules per glucose molecule.^37,38^ This amounts to a high ATP synthesis rate by ATPase and, as a consequence, about 10% of the membrane proteins can be ATPase complexes^39^; indicating that it is one of the top competitors for membrane space. When *E. coli* grows aerobically on other carbon sources than sugars (e.g. on acetate or pyruvate), ATPase is the only source of ATP and is, therefore, an essential enzyme. Also under non-respiratory conditions, ATPase confers a benefit: it can hydrolyze ATP to generate a proton gradient to drive other cellular processes (like membrane transport) and maintain a physiological intracellular pH. ATPase is, therefore, a key enzyme for *E. coli*’s fitness, independent of environmental conditions.

The expression level of ATPase in *E. coli* varies across growth conditions.^37,40–43^ It remains unclear whether these expression levels are optimal or not, i.e. whether they maximize the growth rate across those conditions. We therefore tested ATPase-expression optimality across 26 different carbon sources, each supplemented to M9 minimal medium, and once in complex medium (Luria-Bertani Broth). We exploited a previously characterized *E. coli* mutant (LM3113) and its wildtype, a K12-derived strain (LM3118).^30–32,44,45^ The mutant has an IPTG-titratable *atpIBEFHAGDC* operon. Jensen et al. confirmed that IPTG titration indeed leads to changes in enzyme synthesis.^30–32^ In the 27 different conditions, we grew the IPTG-titratable strain at a range of IPTG concentrations to determine the optimal ATPase-expression level that maximizes the growth rate of the titratable mutant. We also grew the wildtype in this manner to be able to compare its growth rate to that of the mutant. An overview of the experimental approach is shown in Figure 1.

**Figure 1.**
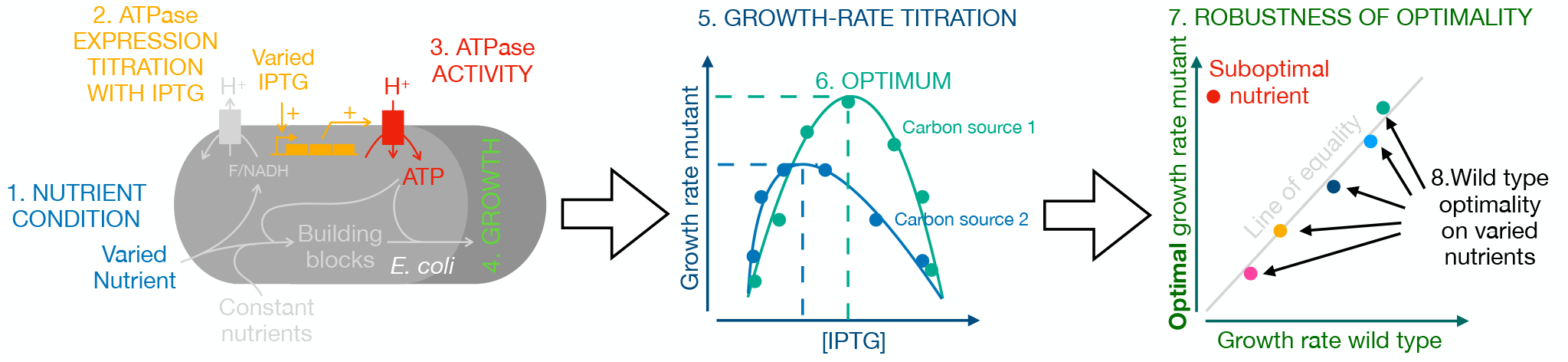
Experimental approach for identification of robust optimal expression of ATPase. *E. coli* wildtype and a ATPase titratable strain were grown on minimal medium in different nutrient conditions. In a single condition, the ATPase concentration was titrated with IPTG and the growth rates of the wildtype and the mutant were measured. The optimal growth rate of the mutant strain was plotted versus the growth rate of the wildtype. If those values lie on the line of optimality for all carbon sources, then *E. coli* displays robust, optimal expression of its ATPase that maximizes the immediate growth rate. Alternatively, for a given nutrient source, the growth rate of the wildtype could be lower than that at the optimal value of the titrated strain, indicating suboptimal expression levels.

All the used nutrients were sole carbon sources, except for: i. arginine, asparagine, glutamine, glycine, and ornithine, which are mixed carbon and nitrogen sources (in those cases, ammonium was not added to the medium) and ii. cytosine, alanine and glucosamine were used as sole nitrogen sources and glucose was added in those cases as the carbon source. The results are shown in Figure 2. The growth-rate titration curves of the wildtype and the mutant are also shown in supplemental figure S1.

**Figure 2.**
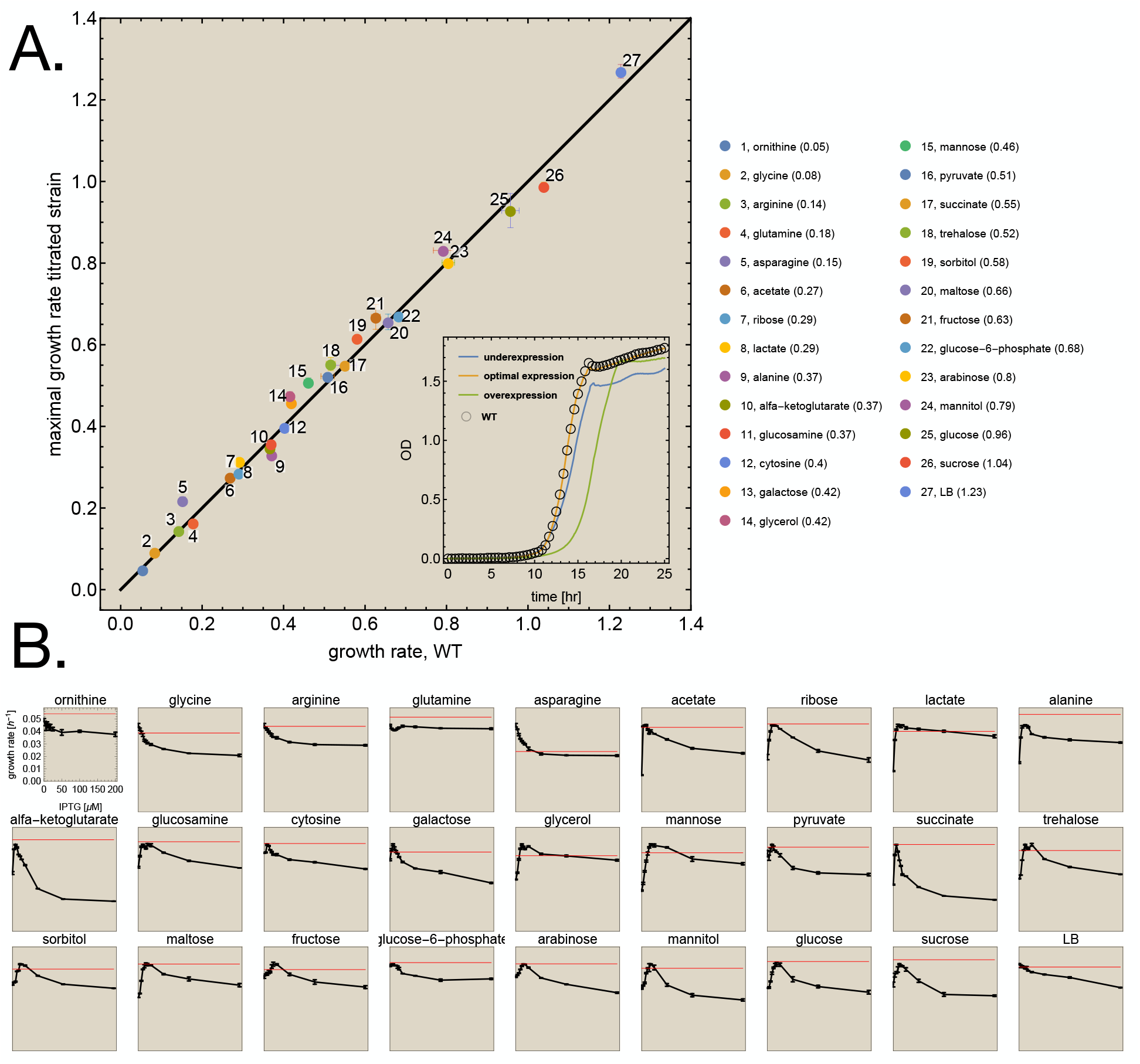
Robust, growth-rate maximizing expression of F_1_F_0_ H^+^-ATPase. The wildtype and the ATPase titratable strain were grown in 27 different conditions. **A**. At each condition, the optimal growth rate of the titratable strain was plotted as function of the growth rate of the wildtype. Error bars indicate the standard deviation. The growth rates of the wildtype and the mutant were compared at the optimal IPTG concentration. The growth rate of the wildtype strain is indicated in the legend after the name of the nutrient and indicated by the red line in the small plots below. The inset shows the growth profiles on glucose of the mutant strain, grown at underexpression, optimal expression and overexpression conditions (as lines), and the wild type as open circles. The profile of the wild type overlaps with that of the mutant at optimal conditions. **B**. For each condition, a plot of the growth rate of the mutant strain (+/- SEM) as function of the IPTG concentration is shown (the axes are only shown once). More detailed plots are shown in the Appendix. The red line indicates the mean growth rate of the wildtype (at the optimal IPTG concentration). Note that the growth rate does not go to zero at high IPTG concentrations. There, the promoter is at maximal activity and ATPase cannot be increased more in concentration. Only with a stronger promoter or multiple gene copies would a higher reduction in growth rate become achievable. We only show the growth rate values and IPTG concentrations for ornithine, the growth-rate range of the other nutrients are different.

Figure 2 indicates that ATPase expression by wildtype *E. coli* is generally very close to optimal behavior. The average absolute deviation from optimality is 6.1±6.1%, indicating that optimal expression of ATPase is surprisingly robust. In a few cases, we found that the wildtype grew faster than the mutant. Note that, due to the experimental setup with a discrete range of IPTG concentrations, the possibility exists that the exact optimal concentration was not included.

A full overview of the growth rates at the optimum and of the wildtype, including the deviation of the wildtype from the optimum can be found in Supplementary Table S1. The absolute deviation values per carbon source are shown in Figure S2. Four carbon sources had a statistically significant deviation (t-test, p-value<0.05) of more than 10%: asparagine (30.3 %), glycerol (12.7 %), alanine (12.5%) and ornithine (10.2%). Of these 4, for ornithine the absolute values of the growth rates only differ by 0.005 h^-1^, and for alanine the wildtype outperforms the apparent optimum (most likely, as stated above, due to the discrete range of IPTG concentrations measured) – pointing towards experimental limitations in these 2 cases. For glycerol, it is interesting to note that an earlier study has also found that the wildtype grows sub-optimally, and can improve after adaptive evolution.^46^ In the case of asparagine, we cannot exclude that the wildtype also shows sub-optimal behavior that could be improved by evolution. When we exclude those four carbon sources, the mean deviation from optimality is 4.4±3.0%. An additional statistical analysis is shown in Figure S3.

Since our main research aim is to get an impression of the robustness of the optimality of expression, we performed a regression analysis to evaluate to what extend the dataset as a whole indicates that the wildtype robustly optimises the expression of ATPase to maximise immediate growth rate. We found a regression coefficient (R^2^) of 0.988, indicating that the wildtype robustly steers ATPase expression to its optimal level throughout all tested conditions.

## Discussion

Studies that focus on growth rate effects of protein expression raise a number of fundamental questions at the interface of microbiology and evolutionary biology.^16,21,22^ Many of them indicate direct^16,19,20,25–30^ or indirect^11,13,15,47,48^ evidence of optimal expression behavior. Together, they suggest that microbial physiology may be predictable and understandable from an evolutionary perspective where selection for maximal growth is constrained by physicochemical and cellular limits. These limits bound cellular protein concentrations and, therefore, the maximally-attainable growth rate.^16^ Optimality of protein expression studies suggest that this optimal state is reached for dozens of key metabolic proteins.^16,19,20,25–30^ In yeast, this was shown for over 70 proteins.^20^ Here we show for the first time that optimal expression of a key metabolic protein can be robust across conditions.

Our findings indicate that the expression of the operon *atpIBEFHAGDC* is optimally regulated by its associated molecular-control system across conditions. Many genes involved in respiration and metabolism are regulated by ArcA and Fnr,^49,50^ but knocking out these proteins did not affect *atp* expression.^37^ Regrettably, how the regulation of the *atp* operon molecularly works is still poorly understood. We note that theoretical studies have shown that optimal gene expression can be achieved by simple circuits composed of basic biochemical interactions, suggesting that even a single transcription factor and a promoter with a single binding site can suffice for optimal steering.^6,9,13,51^ One of the general caveats in our understanding of expression regulation of metabolic genes is that even if we know the identity of the associated transcription factors, we rarely know which metabolites regulate their DNA-binding affinities by binding and stabilizing particular transcription-factor conformations.

Optimal expression of ATPase for immediate growth rate maximization does not necessarily imply that *E. coli* does not carry out a long-term fitness maximization strategy.^6,52^ In fact, evidence exists for long-term fitness maximization too. It has, for instance, been shown that *E. coli* prepares for adverse conditions at slow growth.^53,54^ Nevertheless, a 1% overexpression of an abundant and relatively costly enzyme, like ATPase, represents a significant waste of resources and reduces immediate growth rate. Thus, evolution under constant conditions can be expected to lead to a loss of preparatory overexpression.

We showed here that optimal expression of ATPase is robust across a wide range of nutrient conditions in *Escherichia coli*. This indicates that the expression regulation of a ubiquitous enzyme, central to growth and survival (i.e. fitness), can be understood in terms of growth-rate maximization. Since optimal regulation of gene expression is achievable with simple circuits,^9,13,15,19,51^ optimization of metabolic protein levels for maximal growth rate might be widespread in microbiology. A quite universal principle may exists for regulation of (metabolic) gene expression in microorganism.

## Acknowledgements

We thank Peter Jensen for providing us the *E. coli* strains used in this study. We thank Dirk Bald, Greg Bokinsky, Johan van Heerden, Bas Teusink, Bob Planqué and Hans Westerhoff for in depth discussions.

## Materials & Methods

### Media

For all experiments, M9 minimal medium^55^ was freshly made every experiment day, to avoid degradation. It was supplemented with 1 μg/ml thiamine, as well as the desired carbon source (concentration adjusted to carbon content or nitrogen content; 20 mM for C_6_-carbohydrates, 18.7 mM for N_1_). During preculturing, 10 μM of IPTG was added to the LM3113 strain, to reduce the selective advantage of *tacI* mutants. For the experiments with mixed carbon- and nitrogen sources (arginine, asparagine, glutamine, glycine, ornithine) NH_4_Cl and another carbon source were omitted from the medium; in the cases of cytosine, alanine and glucosamine, NH_4_Cl was omitted but glucose was added to the medium.

### Strains and growth experiments

Derivative strains of *E. coli* K-12 were used in this study, kindly provided by P. R. Jensen. All strains have their lactose permease eliminated, making them reliant on passive diffusion of IPTG. Since lactose permease can act as a costly IPTG-import system its deletion is preferred.

| Strain | Genotype | Phenotype | Reference |
| --- | --- | --- | --- |
| LM3118 | F<sup>+</sup> *asnB32 thi-1 relA1 spoT1 lacUV5 lacY* | *lac* mutant of wildtype | 44,45 |
| LM3113 | As LM3118, but *atpI::(tacIp,lacIOZ)* | IPTG inducible F<sub>1</sub>F<sub>0</sub> H<sup>+</sup>-ATPase | 44,45 |

The obtained strains were sent on LB medium. A single colony was picked and grown on M9 medium+1 µg/ml thiamine+20 mM glucose, then stored in aliquots at -80°C in glycerol stock. For each experiment, an aliquot of the same glycerol stock was used to inoculate the cultures on M9 medium+thiamine+glucose, and grown overnight at 30°C, shaking at 200 RPM. The next morning the OD_600_ was measured, and the cultures propagated to fresh medium containing the desired carbon source at 0.01<OD<0.05, followed by a few more hours of growth (depending on the growth rate) to ensure exponential growth of the cultures.

For the microplate experiments, M9 medium + thiamine + nutrient source was prepared containing a range of IPTG concentrations (0, 3, 6, 9, 12, 15, 18, 25, 50, 100, 200 μM). The OD_600_ of the propagated cultures was measured, and the cells were propagated to the (IPTG containing) media at OD=0.0005, and put in microplate in quadruplicate per treatment. Every microplate also contained LM3118 wildtype on glucose without IPTG as an internal control to confirm proper growth conditions. The spaces in between wells were filled with 0.9% NaCl solution, and the plate sealed at the edges with parafilm to minimize evaporation. The OD_600_ was measured at 30°C every 5 minutes using the Spectramax 384 plus, with shaking in between reads. The data was calibrated based on a dilution series of different ODs measured in a cuvette spectrometer and the microplate reader. The growth rate was then calculated during the exponential growth phase, from the linear ln(OD) range using an R script.

### Statistics

The comparison between the growth rates at the optimal IPTG concentration of wildtype and the titratable strain on all separate carbon sources was tested using a two-tailed, heteroscedastic students t-test. The regression analysis on the overall dataset was done by calculating the total sum of squares 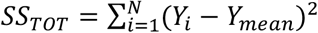 with the growth rate of the wildtype at the optimal [IPTG] on nutrient source *i* for *Y*_*i*_, and the average growth rate of the wildtype throughout conditions for *Y*_*mean*_; the residual sum of squares 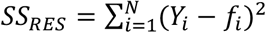 with the growth rate of the titratable strain at the optimum for f_i_; and the coefficient of determination as 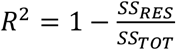

