## Supplemental Information for "*Escherichia coli* robustly expresses ATP synthase at growth rate maximizing concentrations"

Add:

1. Supply all the measured data
2. Supply the Mathematica file for data analysis

| Nutrient source | [IPTG] at optimum | mean $\mu_{max}$ wt | SE $\mu_{max}$ wt | mean $\mu_{max}$ optimum | SE $\mu_{max}$ opt | Deviation wt from opt | t-test (p-value) |
|---|---|---|---|---|---|---|---|
| acetate | 9 | 0.268 | 0.002 | 0.275 | 0.002 | 0.02 | 0.0832 |
| alanine | 12 | 0.371 | 0.002 | 0.330 | 0.002 | -0.13 | 0.0000 |
| alfa-ketoglutarate | 6 | 0.367 | 0.003 | 0.347 | 0.001 | -0.06 | 0.0043 |
| arabinose | 15 | 0.804 | 0.008 | 0.801 | 0.003 | 0.00 | 0.7097 |
| arginine | 0 | 0.143 | 0.000 | 0.145 | 0.003 | 0.01 | 0.5879 |
| asparagine | 0 | 0.152 | 0.003 | 0.218 | 0.004 | 0.30 | 0.0001 |
| cytosine | 6 | 0.403 | 0.002 | 0.397 | 0.003 | -0.01 | 0.2073 |
| fructose | 18 | 0.626 | 0.006 | 0.668 | 0.015 | 0.06 | 0.0642 |
| galactose | 6 | 0.419 | 0.003 | 0.458 | 0.005 | 0.08 | 0.0009 |
| glucosamine | 18 | 0.369 | 0.004 | 0.357 | 0.005 | -0.03 | 0.0886 |
| glucose-6-phosphate | 6 | 0.682 | 0.000 | 0.670 | 0.003 | -0.02 | 0.0317 |
| glucose | 18 | 0.957 | 0.011 | 0.929 | 0.021 | -0.03 | 0.2869 |
| glutamine | 0 | 0.178 | 0.001 | 0.163 | 0.002 | -0.09 | 0.0003 |
| glycerol | 12 | 0.415 | 0.002 | 0.476 | 0.005 | 0.13 | 0.0001 |
| glycine | 0 | 0.084 | 0.002 | 0.092 | 0.002 | 0.09 | 0.0312 |
| lactate | 18 | 0.292 | 0.001 | 0.314 | 0.002 | 0.07 | 0.0001 |
| LB | 0 | 1.228 | 0.005 | 1.270 | 0.008 | 0.03 | 0.0082 |
| maltose | 15 | 0.657 | 0.003 | 0.656 | 0.010 | 0.00 | 0.9399 |
| mannitol | 18 | 0.792 | 0.012 | 0.831 | 0.004 | 0.05 | 0.0412 |
| mannose | 18 | 0.461 | 0.005 | 0.508 | 0.005 | 0.09 | 0.0007 |
| ornithine | 0 | 0.054 | 0.000 | 0.049 | 0.001 | -0.10 | 0.0191 |
| pyruvate | 9 | 0.508 | 0.008 | 0.523 | 0.002 | 0.03 | 0.1723 |
| ribose | 9 | 0.291 | 0.001 | 0.286 | 0.001 | -0.02 | 0.0683 |
| sorbitol | 15 | 0.581 | 0.004 | 0.615 | 0.003 | 0.06 | 0.0004 |
| succinate | 6 | 0.551 | 0.014 | 0.550 | 0.002 | 0.00 | 0.9305 |
| sucrose | 12 | 1.039 | 0.003 | 0.988 | 0.006 | -0.05 | 0.0036 |
| trehalose | 25 | 0.516 | 0.006 | 0.553 | 0.008 | 0.07 | 0.0110 |

**Table S1. Overview for all nutrient sources of wildtype strain (wt) and $F_1F_0$ $H^+$-ATPase titratable strain at the optimum (opt).** Values displayed are the IPTG concentration at the optimum for the titratable strain; the average growth rate ($\mu_{max}$) and standard error (SE) of the wildtype and titratable strain at the optimum; the deviation of the growth rate of the wildtype from the optimum; and the p-value of the corresponding students t-test.

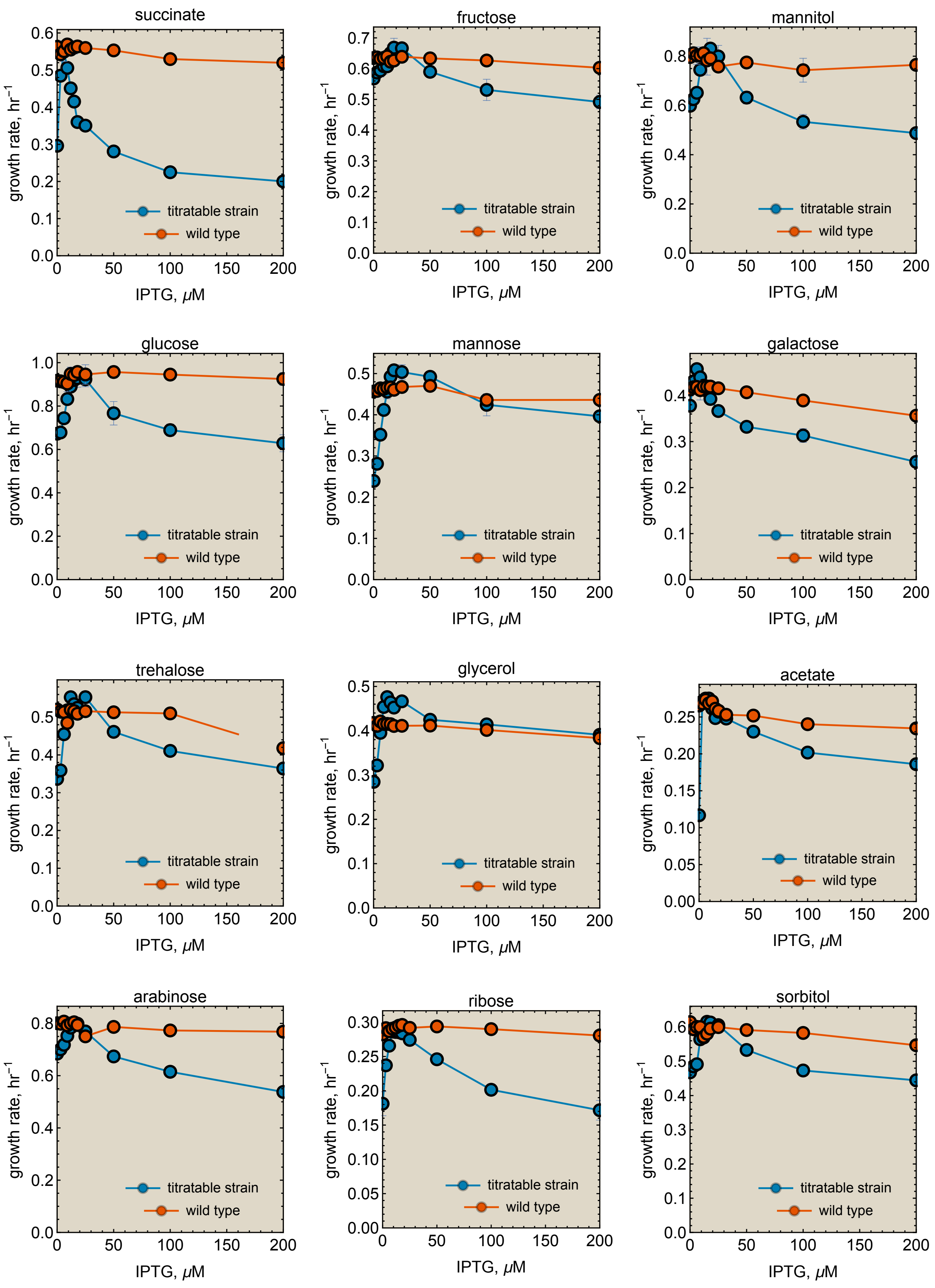


**Figure S1. Growth rate as function of the concentration of IPTG for the $F_1F_0$ $H^+$-ATPase titratable strain and the wildtype (control).** Growth rate +/- std is shown.

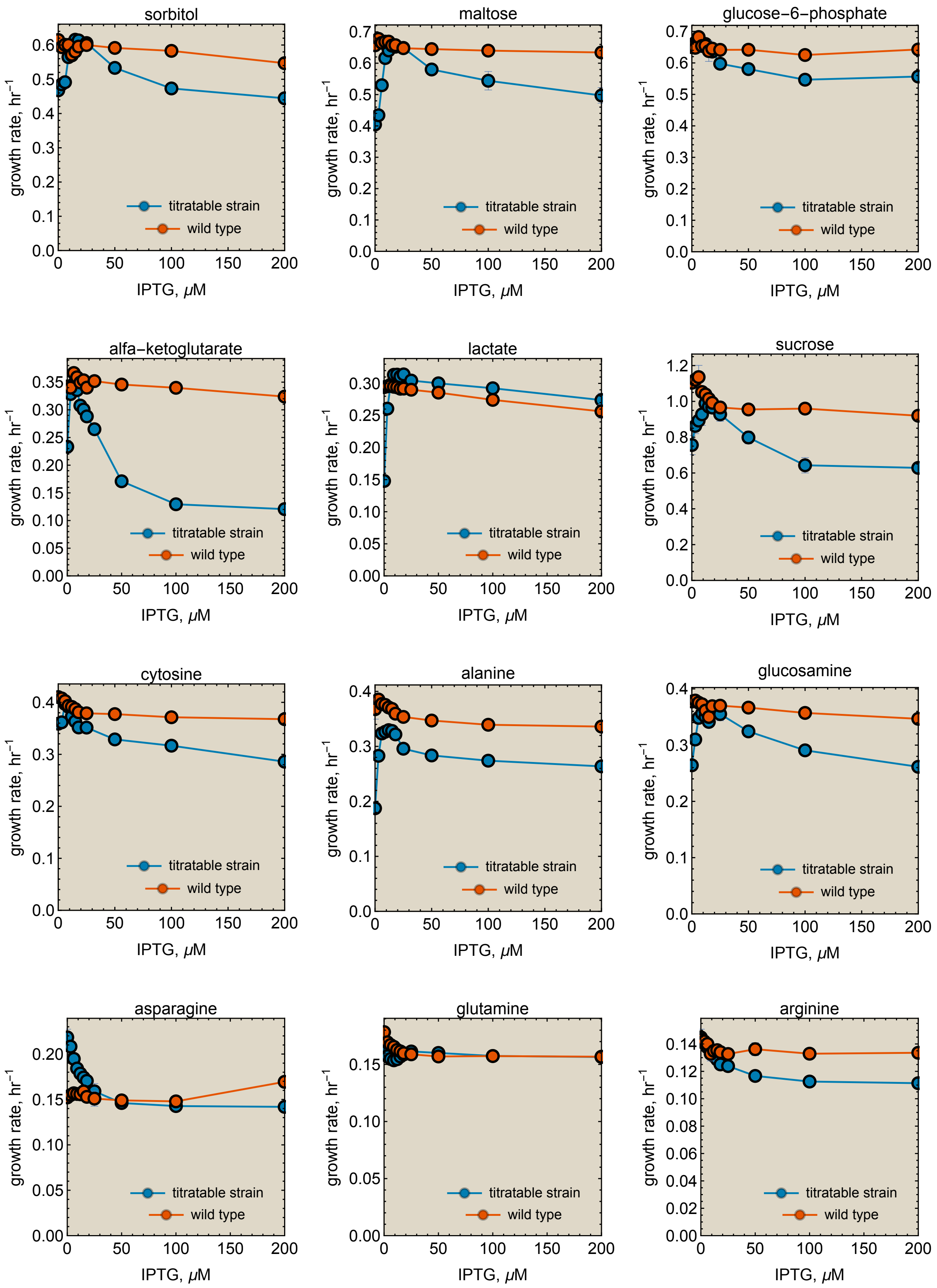


**Figure S1 (continued). Growth rate as function of the concentration of IPTG for the $F_1F_0$ $H^+$-ATPase titratable strain and the wildtype (control).** Growth rate +/- std is shown.

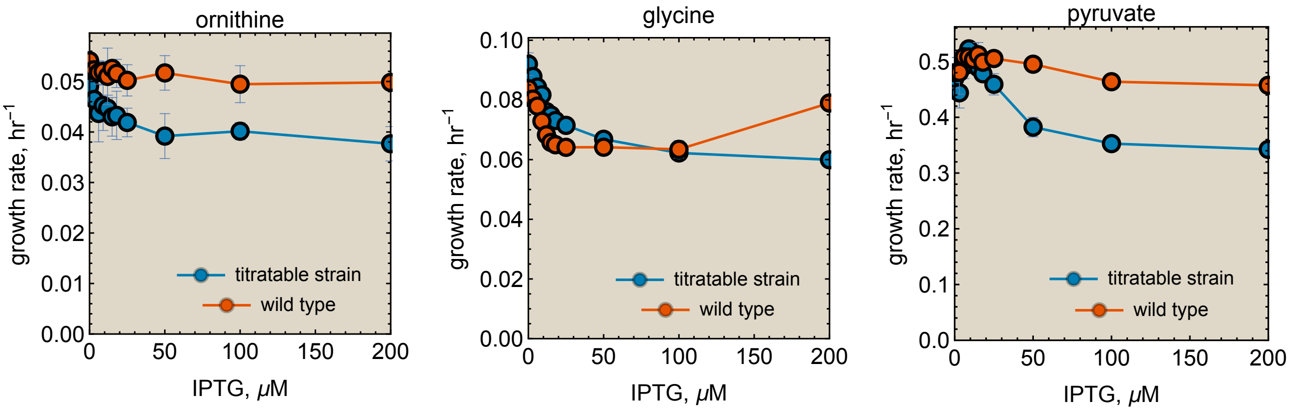


**Figure S1 (continued). Growth rate as function of the concentration of IPTG for the $F_1F_0$ $H^+$-ATPase titratable strain and the wildtype (control).** Growth rate +/- std is shown.

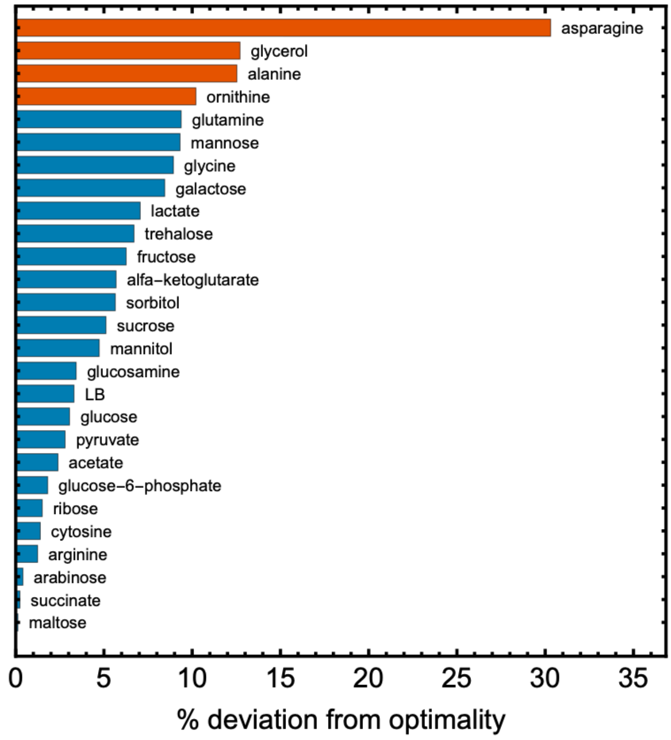


**Figure S2. Percentage absolute deviation from optimal protein expression per carbon source.** Only four carbon sources show a percentage deviation of more than 10%: ornithine (10.2%), glycerol (12.7%), alanine (12.3%) and asparagine (30.3%). Mean absolute deviation is $6.1 \pm 6.1\%$ and with those four carbon sources removed $4.3 \pm 3.0\%$. The growth rates of the wildtype and the mutant were compared at the optimal IPTG concentration.

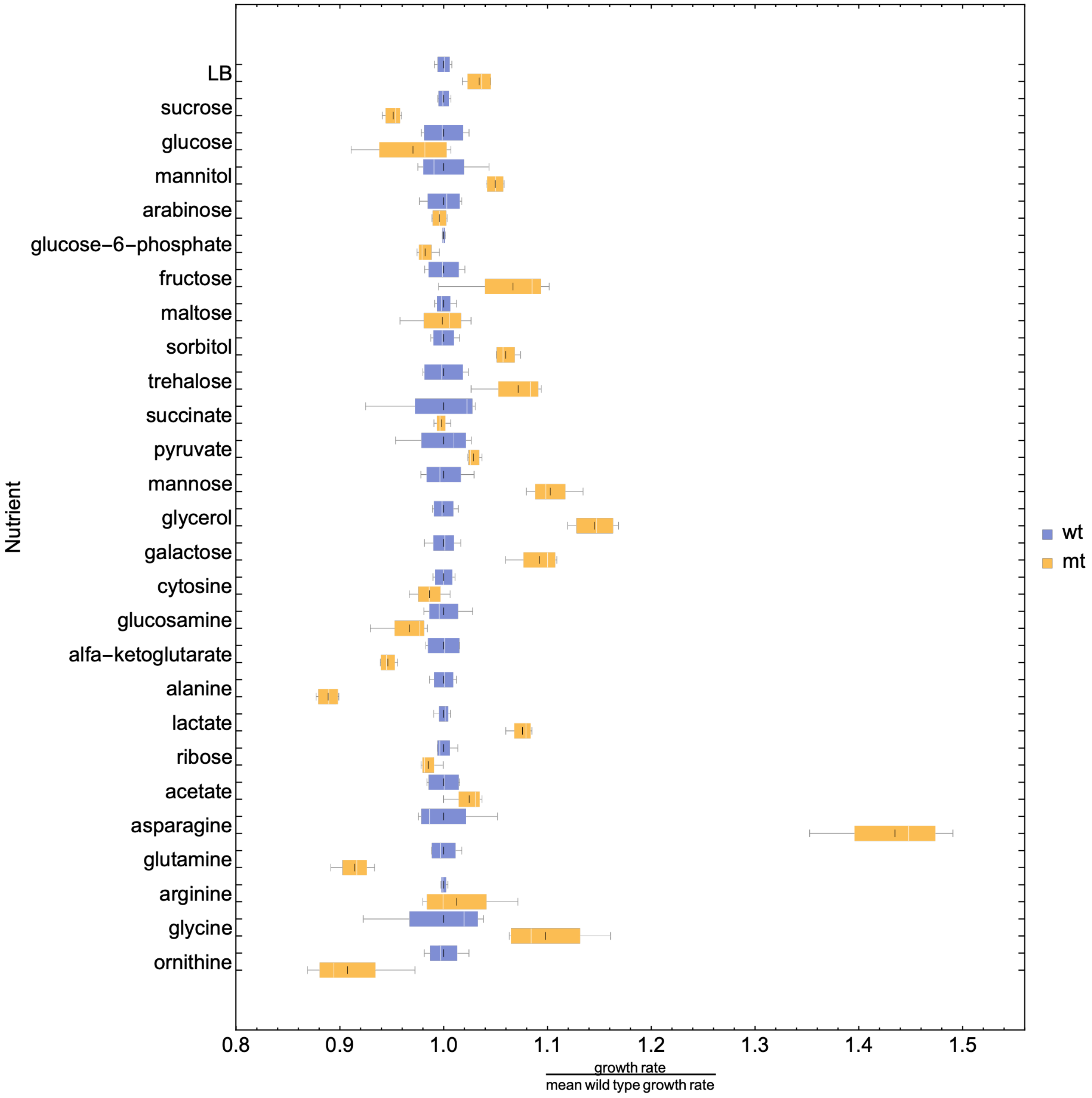


**Figure S3. Box whisker plot of the titratable strain at optimum (mt, yellow) and wildtype (wt, blue) data.** Each nutrient has two boxes, one blue the other yellow, indicating its wild and mutant growth rate values, respectively. The white line indicates the median value, the black line the mean value, the lower end of the box marks the 25% quantile and the upper the 75% quantile.
